# Development of caecaloids to study host-pathogen interactions: new insights into immunoregulatory functions of *Trichuris muris* extracellular vesicles in the caecum

**DOI:** 10.1101/2020.05.11.087684

**Authors:** María A. Duque-Correa, Fernanda Schreiber, Faye H. Rodgers, David Goulding, Sally Forrest, Ruby White, Amy Buck, Richard K. Grencis, Matthew Berriman

## Abstract

The caecum, an intestinal appendage in the junction of the small and large intestines, displays a unique epithelium that serves as an exclusive niche for a range of pathogens including whipworms (*Trichuris spp*). While protocols to grow organoids from small intestine (enteroids) and colon (colonoids) exist, the conditions to culture organoids from the caecum have yet to be described. Here, we report methods to grow, differentiate and characterise mouse adult stem cell-derived caecal organoids, termed caecaloids. We compare the cellular composition of caecaloids to that of enteroids identifying differences in intestinal epithelial cell (IEC) populations that mimic those found in the caecum and small intestine. The remarkable similarity in the IECs composition and spatial conformation of caecaloids and their tissue of origin enables their use as an *in vitro* model to study host interactions with important caecal pathogens. Thus, exploiting this system we investigated the responses of caecal IECs to extracellular vesicles (EVs) secreted/excreted by the intracellular helminth *Trichuris muris.* Our findings reveal novel immunoregulatory effects of whipworm EVs on the caecal epithelium, including the downregulation of responses to nucleic acid recognition and type-I interferon (IFN) signalling.

## 1. INTRODUCTION

The intestine is a continuous tube that stretches from the pylorus to the anus, lined internally by a monolayer of columnar epithelium (Mowat and Agace, 2014). Although continuous, the intestine is composed of defined segments with distinct macro- and microscopic appearances, and specialized functions (Mowat and Agace, 2014; Nguyen et al., 2015). These segments are the duodenum, jejunum and ileum of the small intestine, and caecum, proximal, transverse and distal colon, rectum and anus of the large intestine (Mowat and Agace, 2014; Nguyen et al., 2015).

The caecum is an intestinal appendage at the junction of the small intestine and the large intestine (Burns et al., 2004). This blind-ended sac harbours commensal bacteria that in humans can replenish gut microbiota after disturbances and in the mouse are involved in the fermentative digestion of plant polysaccharides that cannot be digested by enzymes of the small intestine (Al Alam et al., 2012; Backhed et al., 2005; Burns et al., 2004; Eckburg et al., 2005; Mowat and Agace, 2014; Nguyen et al., 2015). Microscopically, the caecum is different from the small intestine because it lacks villi and is more similar to the colon since its mucosa consists of crypts of Lieberkühn with only short regions of flat surface epithelium (Barker, 2014; Mowat and Agace, 2014). Like both the small intestine and colon linings, the caecal epithelium is generated by the division of long-lived intestinal stem cells (ISC) that reside near the bottom of the crypts and produce proliferating transit-amplifying (TA) progenitor cells that later differentiate giving rise to absorptive enterocytes and secretory cells (Paneth, goblet, enteroendocrine and tuft cells) (Barker, 2014). However, the cellular composition of the caecal epithelium is different to that of the small intestine because in the caecum, goblet cells are numerous and found throughout the crypts while Paneth cells are rare (Mowat and Agace, 2014). The colon epithelium presents even larger numbers of goblet cells when compared with the caecum but Paneth cells are absent (Mowat and Agace, 2014; Nguyen et al., 2015). This differential cellular composition contributes to variations in the thickness of the mucus layers overlaying the epithelium and in the microbiota structure (James et al., 2020; McGuckin et al., 2011; Mowat and Agace, 2014). These differences result in distinct niches that are colonised by enteric pathogens, which have successfully evolved to invade and persist in particular intestinal segments.

Understanding the embryonic development of the intestine and the signalling pathways that govern ISC proliferation and differentiation has enabled three-dimensional (3D) organoid cultures to be developed from small intestine and colon adult ISC (Date and Sato, 2015; Sato and Clevers, 2013; Sato et al., 2011; Sato et al., 2009). Organoids are capable of self-renewal and spatial organization, and exhibit similar cellular composition, tissue architecture and organ functionality as their tissue of origin (Date and Sato, 2015; Fatehullah et al., 2016; Li and Izpisua Belmonte, 2019). Culture conditions for enteroids recreate the stem cell niche (SCN), including an extracellular matrix support that mimics the basal membrane component, and a combination of growth factors and morphogens (R-spondin 1, epidermal growth factor (EGF) and Noggin) that stimulate or inhibit the signalling pathways regulating ISC proliferation and differentiation (Date and Sato, 2015; Sato and Clevers, 2013; Sato et al., 2009). A gradient of Wnt signalling, from Paneth cells, is required for the budding of crypt-like structures. The bottom of crypts contain stem and Paneth cells that push proliferating TA cells towards the lumen, where decreasing Wnt levels trigger terminal differentiation of the cells (Sato and Clevers, 2013). Wnt-producing Paneth cells are absent in the colon, so exogenous addition of Wnt ligand (Wnt3A) is required to maintain ISC division in colonoid cultures (Date and Sato, 2015; Sato and Clevers, 2013; Sato et al., 2011). However, the addition of Wnt3A to the medium causes the Wnt gradient to be lost and the organoids to become symmetric round cysts, consisting of a homogeneous population of stem and TA progenitor cells (Sato and Clevers, 2013; Sato et al., 2011). Thus, differentiation of colon organoids into crypt-like structures containing the different epithelial cell lineages requires the withdrawal of Wnt3A (Sato and Clevers, 2013; Sato et al., 2011).

Caecal organoid cultures, hereafter named **caecaloids,** have been generated before using similar culture conditions to those used for colonoids and likewise grow as symmetric round cysts (Miyoshi and Stappenbeck, 2013; Zaborin et al., 2017). However, upon withdrawal of Wnt3A, caecaloids do not recreate the differentiated budding crypt-like structures *(Fig 1A)*. Therefore, an alternative cocktail of growth factors/morphogens is needed to produce caecaloids that showcase the differentiated cells types and 3D spatial organisation present in the caecum.

**Figure 1.**
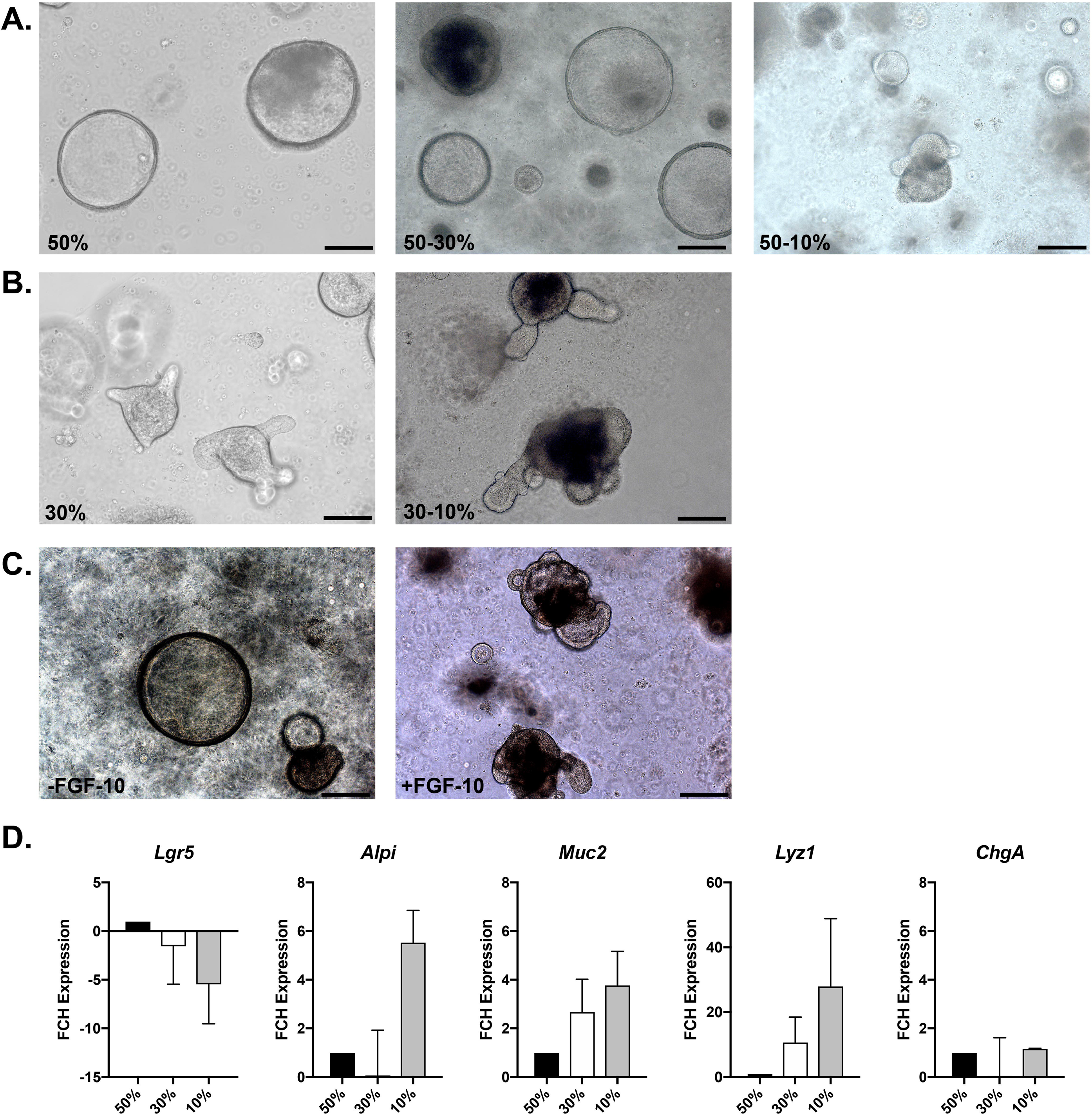
Caecaloid culture and differentiation. Representative bright field microscopy images of caecaloids grown in the presence of 50% (**A**, cystic-undifferentiated morphology) or 30% of Wnt3A-conditioned medium (**B**, budding-differentiated morphology) and differentiated by reduction of concentration to 10%. **(C)** Images of caecaloids grown in 30% of Wnt3A-conditioned medium in the absence (cystic-undifferentiated) or presence (budding-differentiated) of FGF-10. Scale bars 200μm. **(D)** Expression of marker genes measured by qRT-PCR for stem cells (*Lgr5*), enterocytes (*Alpi*), goblet cells (*Muc2*), Paneth cells (*Lyz1*) and enteroendocrine cells *(ChgrA)* in caecaloids grown in 50% and 30% Wnt3A-conditioned medium and further differentiated by reduction of Wnt3A-conditioned medium from 30% to 10%. Results show the mean with standard deviation of results from 2 different caecaloid lines.

The caecal epithelium is the primary colonization site and port of entry for many clinically important pathogens for which mouse models exist, including *Trichuris trichiura (*model organism *T. muris), Salmonella enterica,* serovar Typhymurium, *Campylobacter jejuni, Shigella sonnei, Escherichia coli* (ETEC, model organism *Citrobacter rodentium*), *Yersinia pseudotuberculosis* and *Entamoeba histolytica,* among others (Barthel et al., 2003; Collins et al., 2014; Fahlgren et al., 2014; Houpt et al., 2002; Klementowicz et al., 2012; Lee et al., 1986; Pongpech et al., 1989). Developing mouse caecaloid cultures will enable host interactions of these important pathogens to be studied in an *in vitro* model. Ensuring that these organoids recapitulate the tissue architecture and contain the different IEC types present in the caecum is pivotal to the success of this model system.

Here, we established culture conditions for the long-term expansion, differentiation and characterisation of caecaloid cultures from adult mouse caecal ISC. Caecaloids closely recapitulated the full complement of stem and differentiated cell types present in the caecum, reproducing cellular composition differences between the caecal and small intestinal epithelium. To exemplify the use of caecaloids in the study of host-pathogen interactions in the caecum, we investigated the responses of caecaloids to EVs present in the excretory/secretory (ES) products of the mouse whipworm *T. muris*. EVs are lipid-enclosed structures that can deliver pathogen proteins and nucleic acids into host cells once internalised (Kuipers et al., 2018). *T. muris* EVs can confer protection to whipworm infection in mice (Shears et al., 2018a) and one study has shown *T. muris* EVs are internalised by cells within colonoids (Eichenberger et al., 2018b). Here we examined the functional effects of *T. muris* EVs in caecaloids, which most closely match the *in vivo* context in which the parasites naturally reside. Using RNA sequencing (RNA-seq) of caecaloids microinjected with EVs *T. muris* we discovered a novel immune regulatory function of whipworm EVs on the caecal epithelium, namely the downregulation of responses to nucleic acid recognition and type-I IFN signalling. Our work provides a key tool for future analyses of host interactions with caecal pathogens and their products and identifies new modulatory activities of helminth EVs on IECs.

## 2. MATERIALS AND METHODS

### 2.1 Enteroid and caecaloid culture

Enteroid and caecaloids lines from adult C57BL/6 mice (6-8 weeks old) were derived from small intestinal and caecal epithelial crypts. Briefly, tissues were cut open longitudinally and luminal contents removed. Tissues were then minced, segments were washed with ice cold Dulbecco’s PBS1X without calcium and magnesium (PBS) (Gibco Thermo Fisher Scientific) and vigorous shaking to remove mucus, and treated with Gentle Cell Dissociation Reagent (STEMCELL Tech) for 15 at room temperature (RT) with continuous rocking. Released crypts were collected by centrifugation, washed with ice cold PBS, resuspended in 200 μl of cold Matrigel™ (Corning), plated in 6-well tissue culture plates and overlaid with a Wnt rich medium containing *base growth medium* (Advanced DMEM/F12 with 2mM Glutamine, 10 mM HEPES, 1X penicillin/streptomycin (pen/strep), 1X B27 supplement, 1X N2 supplement (all from Gibco Thermo Fisher Scientific)), 50% Wnt3A conditioned medium (Wnt3A cell line, kindly provided by the Clevers laboratory), 10% R-spondin1 conditioned medium (293T-HA-Rspo1-Fc cell line, Trevigen), 1mM N-acetylcysteine (Sigma-Aldrich), 50 ng/ml rmEGF (Gibco Thermo Fisher Scientific), 100 ng/ml rmNoggin (Peprotech), 10 μM Rho kinase (ROCK) inhibitor (Y-27632) dihydrochloride monohydrate (Sigma-Aldrich), and exclusively for caecaloids, 100 ng/ml rh fibroblast growth factor (FGF)-10 (Peprotech). Organoids were cultured at 37°C, 5% CO_2_. The medium was changed every two days and after one week, pen/strep was take out from the medium and Wnt3A conditioned medium was completely removed for enteroids or reduced to 30% for caecaloids *(expansion medium).* Expanding enteroids and caecaloids were passaged, after recovering from Matrigel using ice-cold PBS or Cell Recovery Solution (Corning), by physical dissociation through vigorous pipetting with a p200 pipette every five to seven days.

For differentiation of caecaloids, after passaging, organoids were grown in *expansion medium* for at least two days to allow reformation and growth in size. Then, medium was changed to *differentiation medium* containing 10% Wnt3A conditioned medium which was replaced every two days for up to four days.

### 2.2 Whole mount immunofluorescence staining of organoids

For whole mount staining, differentiated organoids were recovered from Matrigel using ice-cold PBS, re-plated onto chamber slides (Millicell EZ SLIDE 8-well glass, Millipore) and cultured for additional two days with *differentiation medium*. On the day of staining, organoids were fixed with 4% Formaldehyde, Methanol-free (Thermo Fisher) in PBS for 1 h at RT, washed three times with PBS and permeabilized with 2% Triton X-100 (Sigma-Aldrich) 5% Fetal Bovine Serum (FBS) (Gibco Thermo Fisher Scientific) in PBS for 2 h at RT. Organoids were then incubated with primary antibodies α-villin (1:100, Abcam, ab130751), α-Lysosyme (1:40, Dako, A0099), α-Ki-67 (1:250, Abcam, ab16667), α-Chromogranin A (1:50, Abcam, ab15160), α-Dcamkl-1 (1:200, Abcam, ab31704) and the lectins *Ulex europaeus* agglutinin (UEA, 1:100, Sigma-Aldrich, 19337) and *Sambucus nigra* (SNA, 1:50, Vector Laboratories, FL-1301) diluted in 0,25% Triton X-100 5% FBS in PBS overnight at 15°C. After three washes with PBS, organoids were incubated with secondary antibody (Donkey anti-rabbit 555, 1:400, Molecular Probes, A31572) for 6 h at RT or overnight at 15°C. Organoids were washed three times with PBS and stained with DID (2μg/ml, Biotium, 60014) overnight at 15°C. After three washes with PBS, organoids were counterstained with 4’,6’-diamidino-2-phenylindole (DAPI, 1:1000, Applichem, A1001.0010) at RT for 1 h. Organoids were washed six times with PBS and incubated with FocusClear™ (CelExplorer Labs.) at RT for 1-2 h. Chamber slides were disassembled and mounted using ProLong Gold anti-fade reagent (Life Technologies Thermo Fisher Scientific) and coverslip. Confocal microscopy images were taken with a Leica SP8 and LSM 510 Meta Zeiss confocal microscopes and processed using the Leica Application Suite X (LAS X) software.

### 2.3 Transmission electron microscopy (TEM)

Caecaloids were fixed in 2.5% glutaraldehyde/2% paraformaldehyde in 0.1M sodium cacodylate buffer, post-fixed with 1% osmium tetroxide and mordanted with 1% tannic acid followed by dehydration through an ethanol series (contrasting with uranyl acetate at the 30% stage) and embedding with an Epoxy Resin Kit (all from Sigma-Aldrich). Ultrathin sections cut on a Leica UC6 ultramicrotome were contrasted with uranyl acetate and lead nitrate, and images recorded on a FEI 120 kV Spirit Biotwin microscope on an F416 Tietz CCD camera.

### 2.4 Cell composition analysis of tissues and organoids using ImageStream

Small intestines and caecums of mice were processed individually in parallel. Tissues were open longitudinally, washed with ice cold HBSS 1x (Gibco Thermo Fisher Scientific) containing 1x pen/strep to remove the luminal contents and cut in small fragments. These fragments were incubated at 37°C in DMEM High Glucose (Gibco Thermo Fisher Scientific), 20% FBS, 2% Luria Broth, 1x pen/strep, 100 μg/ml Gentamicin, 10 μM ROCK inhibitor and 0,5 mg/ml Dispase II (Sigma) with horizontal shaking for 90 min to detach epithelial crypts. The crypts containing supernatant were filtered through a 300 μm cell strainer (PluriSelect) and pelleted by centrifugation at 150 g for 5 min at RT. Crypts and enteroids/caecaloids were dissociated into single cells by TrypLE Express (Gibco Thermo Fisher Scientific) digestion 10-20 min at 37°C. The epithelial single-cell suspension was filtered through a 30 μm cell strainer (Sysmex), washed and counted. Cells were fixed with 4% Formaldehyde, Methanol-free in PBS for 20 min at 4°C, washed three times with PBS 1% FBS and permeabilized with 1x Perm/Wash solution (diluted from BD Perm/Wash Buffer 5x in PBS) at RT for 15 min. Cells were then incubated with the primary antibodies used for immunofluorescence staining diluted in 1x Perm/Wash solution for 30 min at 4°C. Cells were washed three times with 1x Perm/Wash solution and stained with secondary antibody/lectins and DAPI diluted in 1x Perm/Wash solution for 30 min at 4°C. After washes with 1x Perm/Wash solution and PBS, cells were resuspended in PBS. Samples were acquired on an Amnis ImageStream MkII Imaging Flow Cytometer (Luminex) at a low speed/high sensitivity flow rate and object magnification at 60x using the INSPIRE software. Data were analysed using the Image Data Exploration and Analysis Software (IDEAS) software. Gating strategy is shown in *Supplementary Figure 3*.

### 2.5 RNA extraction and Quantitative Real-Time PCR (qRT-PCR)

Caecaloids were recovered from Matrigel using Cell Recovery Solution and washed with ice-cold PBS. Caecaloids were then lysed with RTL buffer (RNeasy Mini-kit, QUIAGEN) plus beta-mercaptoethanol (Sigma-Aldrich) and RNA was extracted following manufacturer instructions. Gene expression was quantified by qRT-PCR using ABsolute QPCR Mix, ROX and TaqMan primers (all from Thermo Fisher Scientific) for *Lgr5* (Mm00438890_m1), *Alpi* (Mm01285814_g1), *Muc2* (Mm01276696_m1), *Lyz1* (Mm00657323_m1), *Chga (*Mm00514341_m1) and *Gapdh* (Mm99999915_g1) in a StepOne Real Time PCR system (Applied Biosystems).

### 2.6 T. muris EV purification and quality control

*T. muris* EVs were purified from the ES of *T. muris* as previously described (Shears et al., 2018a). Briefly, the adult parasites were cultured in RPMI medium supplemented with 500 U/ml penicillin and 500 μg/ml streptomycin (Sigma Aldrich) for 18 h (after removing the first 4 hours of ES). The ES was spun at 720 g for 15 min to remove eggs and the supernatant was then filtered using a 0.22 μm filter (Millipore) to further remove debris. Supernatants were then ultracentrifuged at 100,000 g for 2 h in polyallomer tubes using an SV323 rotor. The ultracentrifuge pellet was washed with PBS and re-pelleted by a subsequent spin at 100,000 g for 2 h. The EV pellet was resuspended in 2 ml PBS and stored at −80 prior to further concentration using a 5 kDa MW cut-off vivaspin (Sartorius). The protein content of EVs was quantified using Qubit (Invitrogen) and concentration of EVs quantified by Nanosight (Malvern), resulting in a measurement of 0.18 μg/μl and 2.1×10^7^/μl EVs. EVs were diluted 2:3 with phenol red (Sigma Aldrich) (final concentration of 0.12 μg/μl) prior microinjection into the organoids.

### 2.7 Microinjection of caecaloids

For microinjection, differentiated caecaloids were recovered from Matrigel using ice-cold PBS, re-plated onto microinjection plates (MatTek Corporation) and cultured for additional two days with *differentiation medium*. Microinjections were performed using the Eppendorf TransferMan NK2-FemtoJet express system, in an environmental chamber integrated to a Zeiss Axiovert 200M bright field microscope, to allow all injections to be carried out at 37°C and 5% CO_2_. For RNA sequencing (RNA-seq), 50 caecaloids per microinjection plate were injected with either PBS (as control) or EVs diluted with phenol red so that injected caecaloids could be easily identified. After injection, caecaloids were incubated for 24 h at 37°C, 5% CO_2_, recovered using Cell Recovery Solution and total RNA was extracted as described above.

### 2.8 RNA-seq and analysis

RNA-seq was performed in organoids microinjected with PBS or *T. muris* EVs (n=3) in technical triplicates. Multiplexed cDNA libraries were generated from high-quality RNA samples (RNA integrity number ≥ 7.0) according to the Illumina TruSeq RNA Preparation protocol, and sequenced on an Illumina HiSeq platform. We obtained 3.9 to 4.5 million paired end reads per sample; raw data have been submitted to ENA under the following accessions: ERS2914946, ERS2914953, ERS2914962, ERS2914970, ERS2914978, ERS2914986. Kallisto (v0.43.1)(Bray et al., 2016) was used to pseudoalign reads to the mouse GRCm38 transcriptome (downloaded from Ensembl release 97(Yates et al., 2020), https://www.ensembl.org), with over 92% of reads per sample pseudoaligning. For differential expression analysis, the DESeq function from the DESeq2 package (v1.24.0)(Love et al., 2014) was used to fit a negative binomial GLM for each gene and estimate log2 fold changes, and p values calculated with a Wald test. Genes with an adjusted p value < 0.05 are reported as being differentially expressed. Innate DB v5.4(Breuer et al., 2013) (https://www.innatedb.com) was used to identify enriched Gene Ontology terms. The variance stabilizing transformation function was used to transform counts for Principal Component Analysis (PCA) and heat map plotting. For inclusion in the heatmap, genes were selected based on an association with viral response related GO terms (GO:0051607, GO:0009615, GO:0098586, GO:0039536) by Innate DB and/or Ensembl and an absolute log2fold change > 1.

## 3. RESULTS

### 3.1 Establishment of 3D caecaloid culture conditions

The caecum, like other parts of the intestine, is composed of two layers: 1) an internal endoderm-derived columnar epithelium with absorptive and secretory functions and; 2) an external surrounding mesoderm-derived mesenchyme (Al Alam et al., 2012; Burns et al., 2004). The mouse caecum develops as a bud propagating off the main gut tube early in the differentiation of the gastrointestinal tract (from day 10.5 of embryonic development) (Burns et al., 2004). Epithelial-mesenchymal interactions are critical for the formation of gastrointestinal buds such as the caecum and the stomach. In particular, FGF-10, expressed specifically in the mesenchyme of the caecal bud, signals via the FGF receptor 2b of the epithelium to promote epithelial proliferation at the caecal bud during days 10.5–14.5 of embryonic development (Al Alam et al., 2012; Burns et al., 2004; Zhang et al., 2006). Stomach organoid cultures require FGF-10, in addition to the mouse colonoid culture conditions, to drive budding events and expansion of the cultures (Barker et al., 2010). With this in mind and considering the presence of small numbers of Paneth cells in the caecum (Mowat and Agace, 2014; Nguyen et al., 2015) that contribute Wnt to the culture, we modified existing protocols for colonoid generation (Sato et al., 2011), by adjusting the concentration of Wnt3A in culture medium and including FGF-10.

Upon isolation, crypts were cultured in 50% Wnt3A-conditioned medium until organoids were formed, presenting a cystic morphology *(Fig 1A).* After the first passage, Wnt3A-conditioned medium concentration was reduced to 30% for long-term maintenance culture. At this concentration and in the presence of FGF-10, caecaloids grew as a mixture of cystic and budding organoids and were easily committed to full differentiation by reduction of Wnt3A-conditioned medium concentration to 10% *(Fig 1B* and *C)*. FGF-10 was critical to drive budding events on caecaloids *(Fig 1C)*, just as in stomach organoids (Barker et al., 2010). In the absence of Wnt3A (enteroid culture condition) caecaloids did not survive, indicating the requirement of Wnt3A addition to the medium for their expansion (data not shown). However, when long-term culturing caecaloids with 50% Wnt3A-conditioned medium, as required for the culture of colonoids, it was not possible to induce their differentiation by withdrawal of Wnt3A *(Fig 1A)*. These results indicate caecal ISC have specific growth factor requirements for division and differentiation, which are modelled *in vitro* by fine-tuning the addition of exogenous Wnt ligands and mimicking epithelial-mesenchymal interactions by addition of FGF-10.

### 3.2 Differentiated caecaloids closely recapitulate the caecal epithelium

While budding morphology is a sign of differentiation of organoids, we next sought to evaluate if differentiation culture conditions (initial culture after passaging in expansion medium with 30% Wnt3A-conditioned medium for 2 days, followed by 4 days culture in differentiation medium with 10% Wnt3A-conditioned medium) resulted in full epithelial maturation of caecaloids recreating the caecal epithelium. Therefore, to characterise the caecaloid cellular composition, we first used qRT-PCR to evaluate the expression of known IEC populations markers, including Leucine-rich repeat-containing G-protein coupled receptor 5 (*Lgr5)* for stem cells, alkaline phosphatase (*Alpi*) for absorptive enterocytes, mucin 2 (*Muc2*) for goblet cells, lysozyme 1 (*Lyz1*) for Paneth cells and chromogranin A (*ChgA*) for enteroendocrine cells *(Fig 1D)*. Gene expression was measured in caecaloids maintained with 50% or 30% Wnt3A-conditioned medium and in differentiated caecaloids cultured as described above. All markers were detected confirming the presence of all cellular populations. When comparing caecaloids grown in the presence of 50% and 30% Wnt3A-conditioned medium, we observed a downregulation of *Lgr5* in the latter group indicating a reduction in the number of stem cells *(Fig 1D).* Conversely, growing organoids under expansion conditions (30% Wnt3A) induced the expression of *Muc2* and *Lyz1*, suggesting an increase in goblet and Paneth cells, respectively *(Fig 1D)*. Upon differentiation of caecaloids by decreasing Wnt3A-conditioned medium concentration to 10%, we observed further downregulation of *Lgr5* and detected upregulation of *Alpi* indicating an increase in absorptive enterocytes *(Fig 1D)*. These results agree support our morphological observations *(Fig 1B)* of further differentiation of organoids upon reduction of the concentration of Wnt3A in the culture medium *(Fig 1B)*.

To further study the differentiation status of the caecaloids we performed confocal immunofluorescence microscopy of caecaloids and enteroids *(Figs 2* and *3).*The cellular markers analysed were Ki-67 (present in proliferating stem and TA cells), villin (staining microvilli on absorptive enterocytes), chromogranin A (marker of enteroendocrine cells), lysozyme (produced by Paneth cells), Dclk-1 (identifying tuft cells), and a combination of the lectins UEA and SNA that bind mucus on goblet cells. We observed that differentiated caecaloids contained the following: proliferating cells (Ki-67^+^) at the bottom of budding regions *(Fig 2A),* microvilli of enterocytes bordering the lumen *(Fig 2B),* few enteroendocrine (*Fig 2C*) and tuft cells *(Fig 2D)* but numerous goblet cells *(Fig 2A-D)*. The presence of enterocytes, goblet and enteroendocrine cells in caecaloids was confirmed using TEM *(Fig 2E)*. We did not detect Paneth cells in caecaloids using this methodology *(Supplementary Fig 1)*. In contrast, enteroids have numerous Paneth cells *(Fig 3E)* present at the bottom of budding regions where Ki-67+ cells are also located. Enteroids have fewer goblets cells *(Fig 3A-E)*, and show similar levels of enteroendocrine *(Fig 3C)* and tuft cells *(Fig 3D)* when compared with caecaloids *(Fig 2)*.

**Figure 2.**
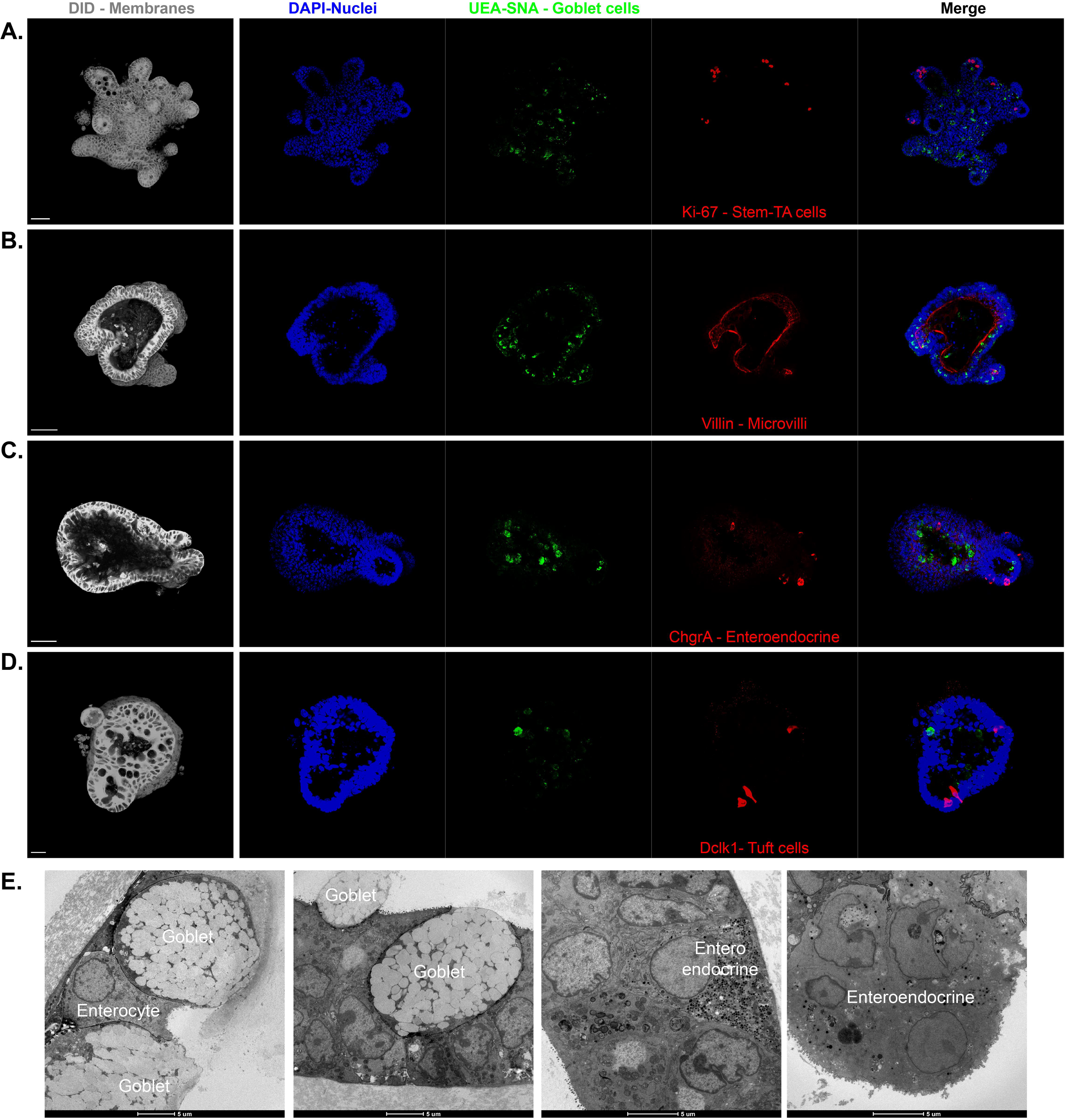
Confocal microscopy characterisation of differentiated caecaloids. Images of caecaloids expanded and further differentiated by reduction of Wnt3A-conditioned medium from 30% to 10%, showing the presence of all IEC populations. **A-D.** Confocal IF microscopy with antibodies staining **(A)** Ki-67, marker of proliferating cells, stem and TA cells; **(B)** Villin, identifying microvilli of enterocytes; **(C)** Chromogranin A expressing enteroendocrine cells; **(D)** Dclk-1, marker of tuft cells; and with the lectins UEA and SNA that bind mucus in goblet cells. DAPI stains nuclei and DiD the cell membranes. Scale bar 50μm for A, B, C and E, 20μm for D. **G.** TEM images showing enterocytes, goblet and enteroendocrine cells present in caecaloids.

**Figure 3.**
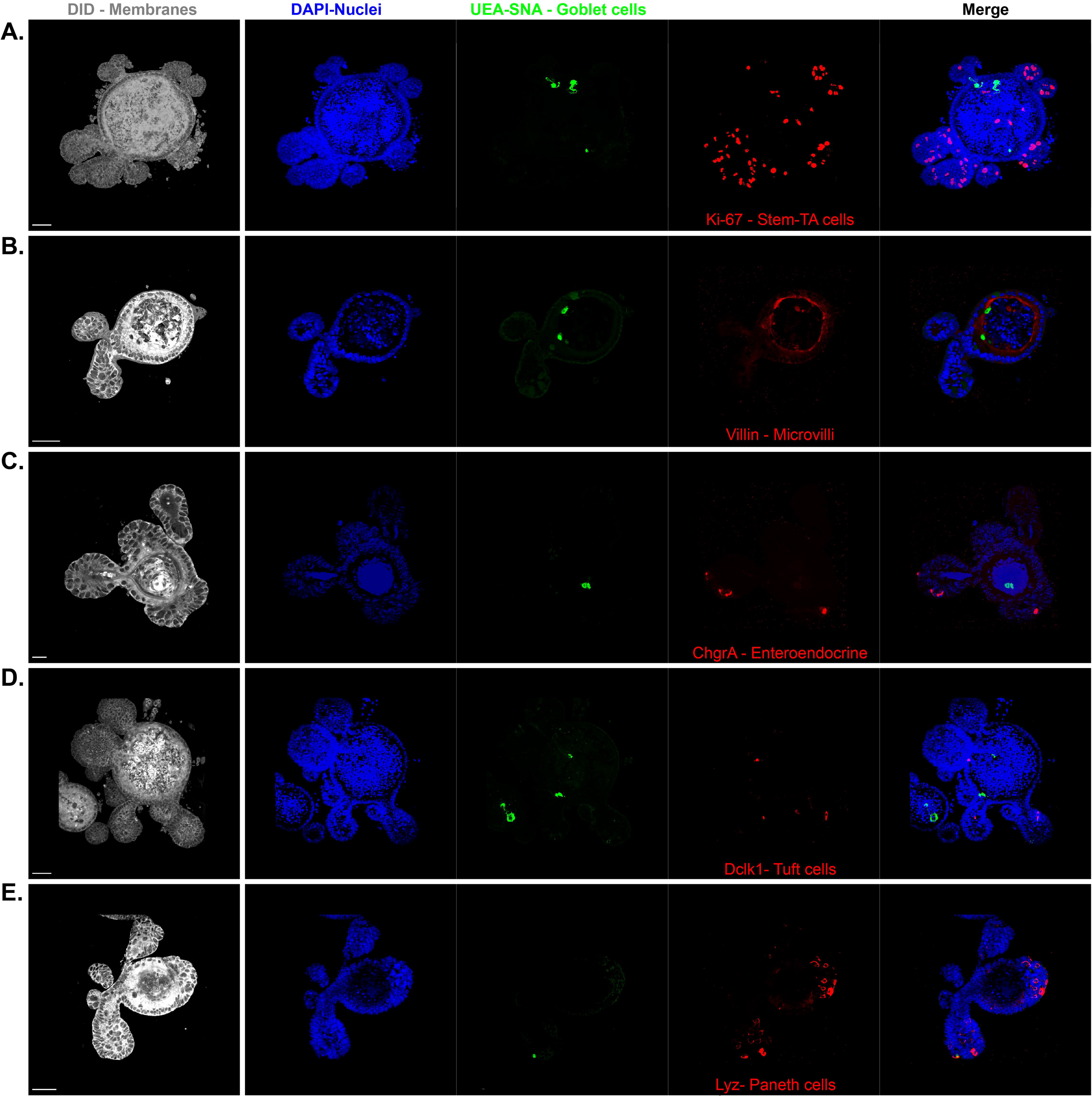
Confocal microscopy characterisation enteroids. Confocal IF microscopy images of enteroids, showing the presence of all IEC populations stained with antibodies for **(A)** Ki-67, marker of proliferating cells, stem and TA cells; **(B)** Villin, identifying microvilli of enterocytes; **(C)** Chromogranin A expressing enteroendocrine cells; **(D)** Dclk-1, marker of tuft cells; **(E)** Lysozyme expressing Paneth cells; and with the lectins UEA and SNA that bind mucus in goblet cells. DAPI stains nuclei and DiD the cell membranes. Scale bar 50μm for A, B, D and E, 20μm for C.

Next, we aimed to determine how caecaloids and enteroids reflect the cellular composition of the tissue of origin. Thus, to quantitatively characterise the different cellular populations in tissue and organoids, we performed ImageStream analysis on single cell preparations stained with antibodies and lectins for markers of the major IECs populations. ImageStream combines both bright field and fluorescence microscopy, coupled with flow cytometry capabilities, allowing enterocytes as well as stem, enteroendocrine, Paneth and goblet cells to be clearly identified and quantified *(Fig 4A* and *Supplementary Fig 2)*. We found a remarkable similarity in the percentages of the different cell types in the organoids and the tissue from which they were derived *(Fig 4B)*. Moreover, these results confirmed our observations using confocal immunofluorescence staining that showed proportionally more goblet cells in the caecal tissue and caecaloids than in the small intestine and in enteroids *(Fig 4B)*. Conversely, the proportions of Paneth cells are lower in the caecum and caecaloids when compared with small intestine and enteroids *(Fig 4B)*. The proportions of enterocytes, enteroendrocrine and stem cells are similar among both tissues and organoids *(Fig 4B)*. Together these data demonstrate that our methods allow the generation of caecaloids closely recapitulating the cellular composition and architecture of the caecal epithelium.

**Figure 4.**
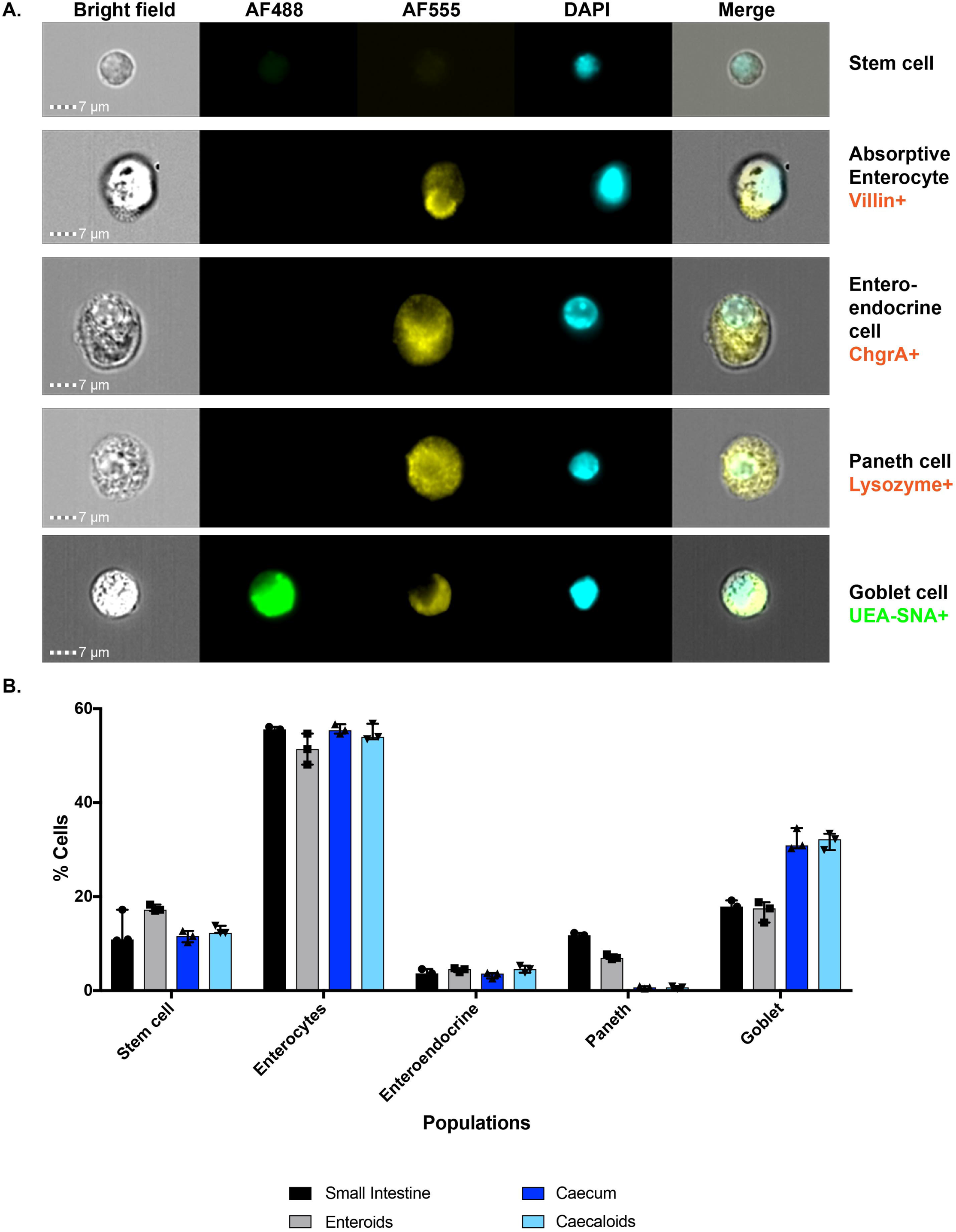
Comparison of cellular composition of enteroids, caecaloids and small intestine and caecum tissues by ImageStream. Enteroids, caecaloids and IECs from small intestine and caecum were dissociated into single cells, stained with antibodies and lectins targeting enterocytes, enteroendocrine, Paneth and goblet cells and visualized by ImageStream. **A.** Bright field and fluorescence representative images of cellular populations. Scale bar 7μm. **B.** Percentages (median with interquartile range) of cellular populations identified by ImageStream. n=3. Note the consistency on the composition of organoids and the tissue of origin.

### 3.3 Caecaloids as a model to study host-pathogen interactions: understanding the effects of *T. muris* EVs on caecal IECs

Next, we aimed to use caecaloids as an *in vitro* model to study interactions between pathogens invading the caecum and the epithelium of this organ. Whipworms are intracellular helminths that inhabit the caecal epithelium. In order to persist in their host, whipworms modulate intestinal inflammation ensuring their host and their own survival (Grencis, 2015; Klementowicz et al., 2012). One such mechanism of immunomodulation is the release of ES products that can interact with immune cells, the regulatory impact of which has been described (Bancroft et al., 2019; Klaver et al., 2013; Kuijk et al., 2012; Laan et al., 2017; Leroux et al., 2018; Shears et al., 2018b). ES products likely act also on caecal IECs given whipworms live inside the epithelium; however, little is known about these interactions (Hiemstra et al., 2014). One component of whipworm ES are EVs, lipid membrane enclosed structures with the capacity to transfer a multitude of nucleic acids and proteins to a single cell at once, and that have been shown to be potent host modulators (Buck et al., 2014; Coakley et al., 2017; Eichenberger et al., 2018a; Eichenberger et al., 2018b; Hansen et al., 2015; Shears et al., 2018a; Tritten et al., 2017). To date, the responses of caecal IECs to whipworm EVs have not been studied. To investigate these interactions, we purified EVs from the ES of *T. muris* adult worms *(Supplementary Fig 2)* and microinjected the EVs into caecaloids. As the apical surface of the IECs in 3D caecaloids is facing the lumen *(see microvilli (villin) staining Fig 2B)*, microinjection is therefore required to mimic the interactions that naturally take place in the caecum (Duque-Correa et al., 2020). PBS was microinjected in 3D caecaloids as a control. After 24 h of culture, total RNA was extracted and gene expression changes in response to EV administration were evaluated by RNA-seq. We observed a clear response of the caecaloids to the EVs *(Fig 5A* and *B)* with a total of 88 genes upregulated and 173 genes downregulated. Interestingly, stimulation with EVs secreted by adult *T. muris* parasites resulted in significantly reduced expression of viral response associated genes by caecal IECs *(Fig 5C)*. Specifically, we detected decreased expression of genes involved in the cytosolic sensing of nucleic acids including *Dhx58, Ddx60* and *Irf7*, which are part of the signalling cascade that results upon engagement of retinoic-acid inducible gene I (RIG-I)-like receptors by dsRNA (Liu et al., 2016). Consequently, EVs treatment of caecaloids resulted in downregulation of interferon stimulated genes (ISGs), comprising *Oasl2*, *Oas2* and *3, Ifit1* and *3*, *and Isg15,* which are transcribed in response to nucleic acid recognition and type-I IFN signalling (Perng and Lenschow, 2018). Although this is just one example of the downstream use of caecaloids, our results suggest the anti-inflammatory effects of whipworm infections and their ES products can be, at least in part, mediated by a direct effect on the caecal epithelium.

**Figure 5.**
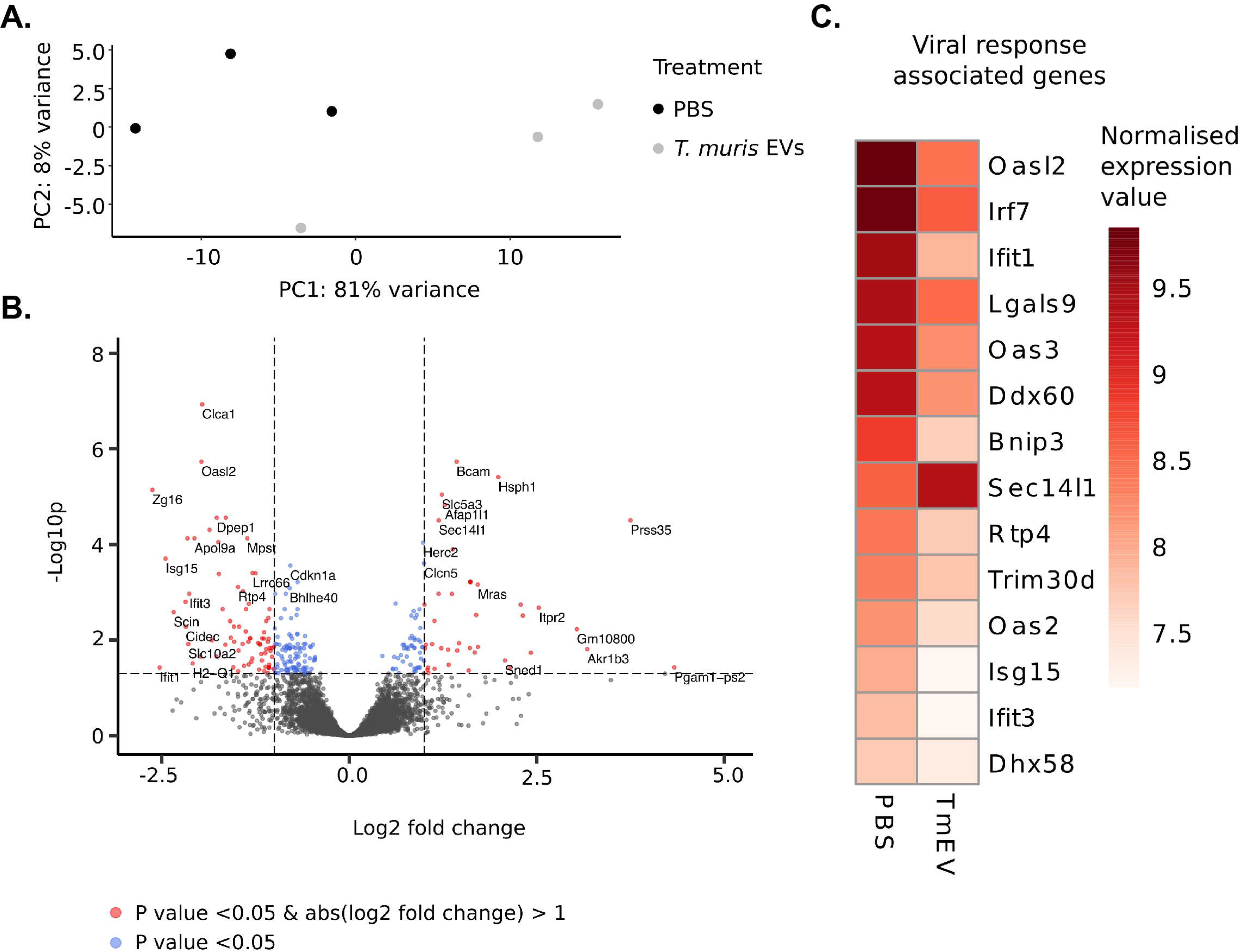
Transcriptional response of caecaloids to microinjection of *T. muris* EVs. **A.** Principal component (PC) analysis showing sample clustering across PC1 and PC2. **B.** Volcano plot showing transcriptional response to microinjection of *T. muris* EVs. Genes significantly differentially expressed (adjusted p value < 0.05) are indicated in red (absolute log2 fold change > 1) or blue (absolute log2 fold change < 1). **C.** Heat map of transformed and normalised expression counts for selected genes (mean of three replicates is represented). Viral response associated genes are associated with the GO terms GO:0051607 and/or GO:0009615 by Innate DB (https://www.innatedb.com/) and/or Ensembl (https://www.ensembl.org).

## 4. DISCUSSION

Here we developed culture conditions for the long-term maintenance and differentiation of caecaloids closely resembling the composition and spatial conformation of the caecal epithelium. Our methods fill a crucial gap on the protocols to generate organoids recreating the differences on the epithelium that distinguish all intestinal segments and that are crucial in the use of these *in vitro* systems to model host-enteric pathogen interactions.

Several pathogens have a tropism for the caecum and the particularities of its mucosa provide a defined niche for which these bacteria and parasites have evolved mechanisms to invade and colonise. In particular, whipworms are large metazoan parasites that live in the caecum of their host, where they tunnel inside IECs creating a multi-intracellular niche (Klementowicz et al., 2012; Tilney et al., 2005). Whipworms can remain in their host for years causing chronic infections. To optimise their residence in their hosts, whipworms manipulate host inflammation partly through the immunoregulatory effects of ES products released by the parasites (Bancroft et al., 2019; Eichenberger et al., 2018b; Hansen et al., 2015; Klaver et al., 2013; Kuijk et al., 2012; Laan et al., 2017; Leroux et al., 2018; Shears et al., 2018a; Shears et al., 2018b; Tritten et al., 2017). Progress has been made to understand the composition and anti-inflammatory actions of whipworm ES products. This has largely involved proteomic analyses of the ES products and, more recently, characterisation of the protein and nucleic acid cargo of EVs (Eichenberger et al., 2018b; Hansen et al., 2015; Leroux et al., 2018; Shears et al., 2018a; Shears et al., 2018b; Tritten et al., 2017; White, 2020). Moreover, immunomodulatory effects of adult *T. suis* (natural pig whipworm) and *T. muris* ES products on different immune cells have been described (Bancroft et al., 2019; Klaver et al., 2013; Kuijk et al., 2012; Laan et al., 2017; Leroux et al., 2018). In contrast, very little is understood regarding the modulatory functions of ES products, or EVs in particular, on IECs, with only one report showing that *T. suis* ES stimulation of an epithelial cell line results in reduced barrier function and decreased lipopolysaccharide-induced TNF-α and CXCL1 production (Hiemstra et al., 2014). IECs are important sensors of intestinal helminth infections initiating innate immune responses and with specialised effector functions that contribute to the expulsion (Artis and Grencis, 2008; Grencis, 2015). Compared to cell lines, organoids more accurately reproduce the composition and architecture of the intestinal epithelium and recently have started to be used to characterise the interactions and IECs responses to ES products of various helminths (Duque-Correa et al., 2019). Specifically, stimulation of murine enteroids with *Trichinella spiralis* ES products and extracts indicated that sensing of parasitic products by tuft-cell receptors results in Ca^2+^ responses (Luo et al., 2019). Moreover, imaging experiments of murine enteroids and colonoids microinjected with EVs present in the ES of *Nippostrongylus. brasiliensis* and *T. muris,* respectively, showed their uptake by host IECs (Eichenberger et al., 2018a; Eichenberger et al., 2018b). A similar approach has been used to visualize the internalization of *Ascaris suum* EVs co-cultured with canine enteroids (Chandra et al., 2019). However, to date organoids have not been exploited to study host IECs responses to helminth EVs.

Here, for the first time, we evaluated the functional effects of helminth EVs on IECs using organoids. Microinjection of adult *T. muris* EVs in fully differentiated caecaloids resulted in downregulation of expression of viral response associated genes by caecal IECs, including those involved in cytosolic sensing of nucleic acids via RIG-I-like receptors and ISGs produced in response to nucleic acid recognition and type-I IFN signalling (Liu et al., 2016; Perng and Lenschow, 2018). Intriguingly, *T. muris* EVs contain parasite small RNAs (sRNAs) (Eichenberger et al., 2018b; Tritten et al., 2017; White, 2020), which instead of triggering host responses to foreign RNA appear to supress such detection mechanisms. This may enable EVs functions, allowing foreign RNA cargo to operate without being sensed by cell. Recent publications have shown type-I IFN responses are induced in response to helminth infections. Particularly, stimulation with *Schistosoma mansoni* antigens (Webb et al., 2017) and infection with *N. brasiliensis* (Connor et al., 2017) results in type-I IFN signalling in dendritic cells, which is required for initiation of Th2 responses. In the setting of *Heligmosomoides polygyrus* infection, type-I IFN responses are reported to be upregulated in the duodenum (McFarlane et al., 2017) and inhibit granuloma formation around larval parasites (Reynolds et al., 2014). Interestingly, type-I IFN responses have not been previously associated to whipworm infections, but their relevance on the development of type 2 immunity in other helminths suggest that by blocking them adult whipworms may counteract host immune responses that result in their expulsion. Our findings open therefore a new avenue of investigation on the interactions of the worm with its host cells and the role of IECs as sensors and orchestrators of the immune responses against whipworms (Artis and Grencis, 2008). In the near future, we aim to understand the mechanisms by which the nucleic acids and protein cargo of the EVs exert such functions in caecaloids. In this regard, our studies on the sRNAs composition of *T. muris* EVs presented on this special issue (White, 2020) will be critical in the identification of targets in the host IECs. Moreover, our methods for microscopy characterisation of IECs populations in caecaloids will be pivotal to pinpoint IEC types preferentially internalising EVs and their intracellular interactions. These future experiments will also shed light into the immunoregulatory effects of therapies using live parasitic worms including whipworms *(T. trichiura* and *T. suis)*, worm secretions and worm-derived synthetic molecules that are being trialled to treat Intestinal Bowel Diseases (IBD) (Smallwood et al., 2017; Varyani et al., 2017).

The remarkable recapitulation of the caecal epithelium achieved by caecaloids will similarly allow the interactions of other caecal pathogens and commensals with the IECs of this organ to be studied with precision. In particular, the multicellularity of this *in vitro* system can be exploited to investigate the role of different IECs populations in pathogen invasion and colonization, host damage and responses (Duque-Correa et al., 2019). The up- and down-regulation of cell populations and factors in caecal-specific context can also be evaluated after exposure to pathogens and their products (Duque-Correa et al., 2019). In addition, caecaloids can be used in studies investigating how caecal microbiota impact the caecal epithelium composition and metabolism. Moreover, caecaloids could be used to model inflammatory pathologies of the caecum, including cancer and IBD, and better understand their aetiology and compare it with inflammation present in other intestinal segments.

In the future, complementation of caecaloid cultures with other tissue components including cellular populations of the SCN (stromal and immune cells), commensal microbiota, chemical gradients and physical/mechanical forces (Barrila et al., 2018; Duque-Correa et al., 2019; Fatehullah et al., 2016; Takebe and Wells, 2019) will more closely recreate the caecal native microenvironment and provide a more complex model to investigate caecal pathologies and the intricacies of pathogen mechanisms to colonise and modulate these niches.

## ACKNOWLEDGEMENTS

This work was supported by the National Centre for the Replacement, Refinement and Reduction of Animals in Research (UK) David Sainsbury Fellowship Grant NC/P001521/1; the Rosetrees Trust (UK) Grant M813; the Wellcome Trust (UK) Grants Z10661/Z/18/Z/WT and Z03128/Z/16/Z/WT; and The Royal Society (UK) Grant IC160132.

## SUPPLEMENTARY FIGURE LEGENDS

**Supplementary Figure 1.**
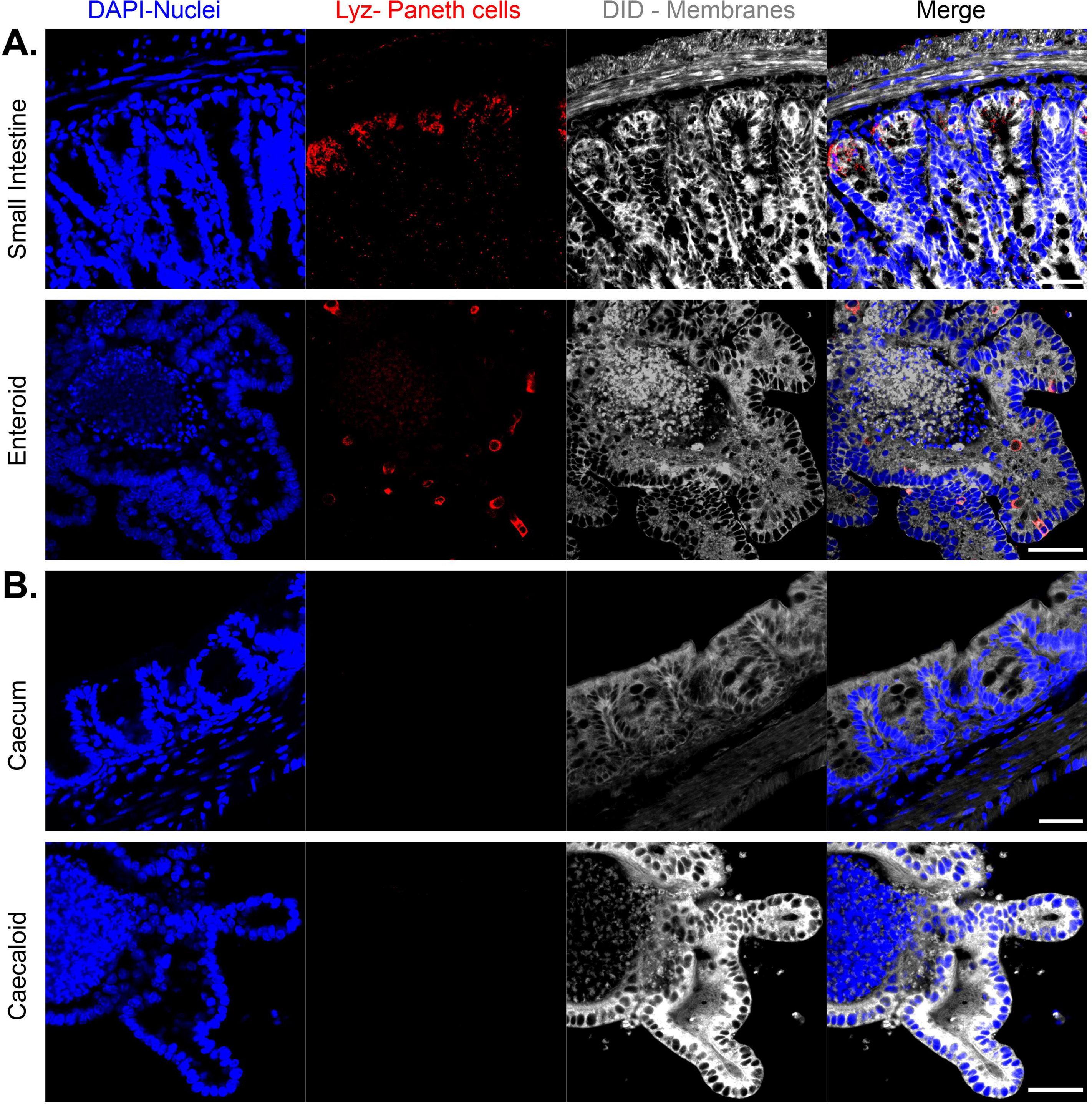
Lysozyme staining as a marker of Paneth cells in small intestine and caecum tissue and organoids. Images of confocal IF microscopy with an antibody staining Lysozyme expressing Paneth cells in small intestine tissue and organoids **(A),** which are not frequently found in caecal tissue and organoids **(B)**. DAPI stains nuclei and DiD the cell membranes. Scale bar 100μm.

**Supplementary Figure 2.**
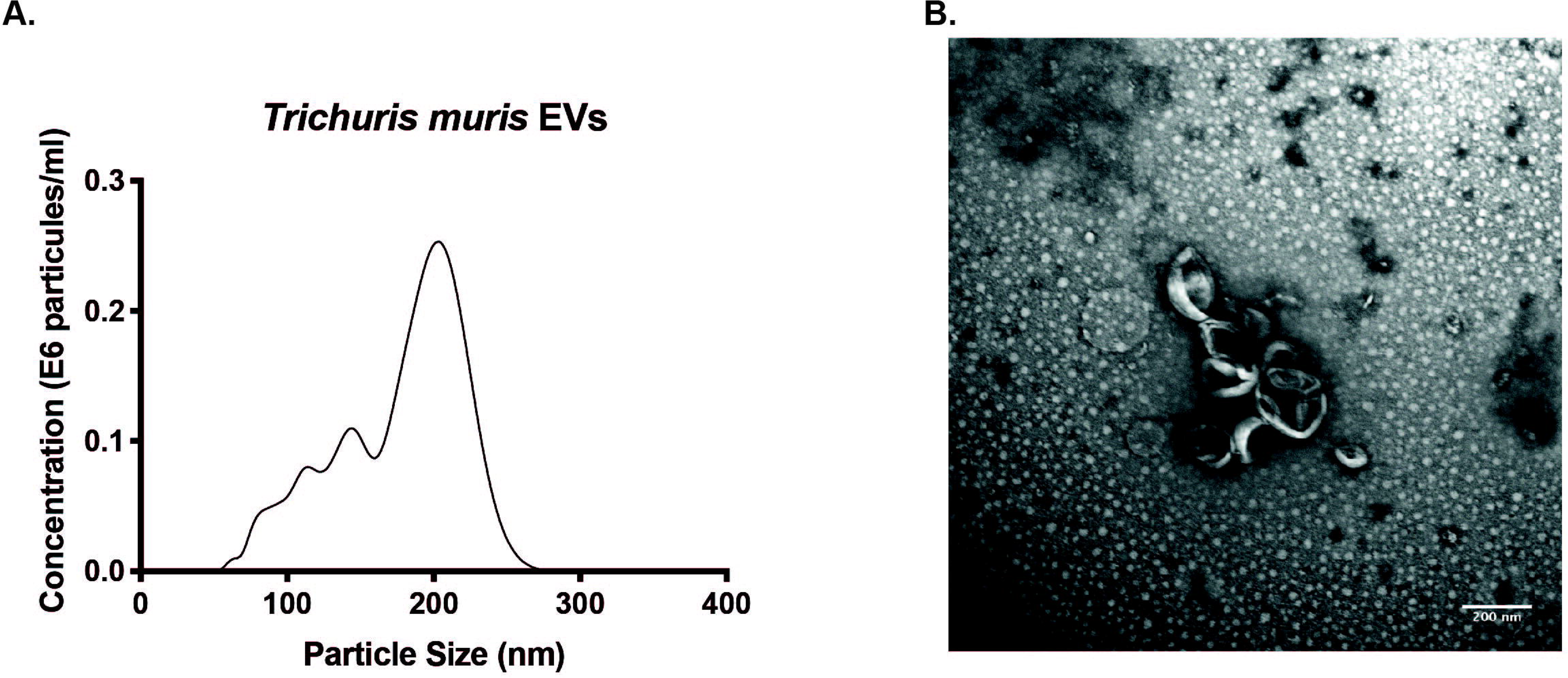
**Qualitative analysis of *T. muris* EVs** by **(A)** Transmission EM and **(B)** Profile of EVs by Nanosight, at 1:1000 dilution.

**Supplementary Figure 3.**
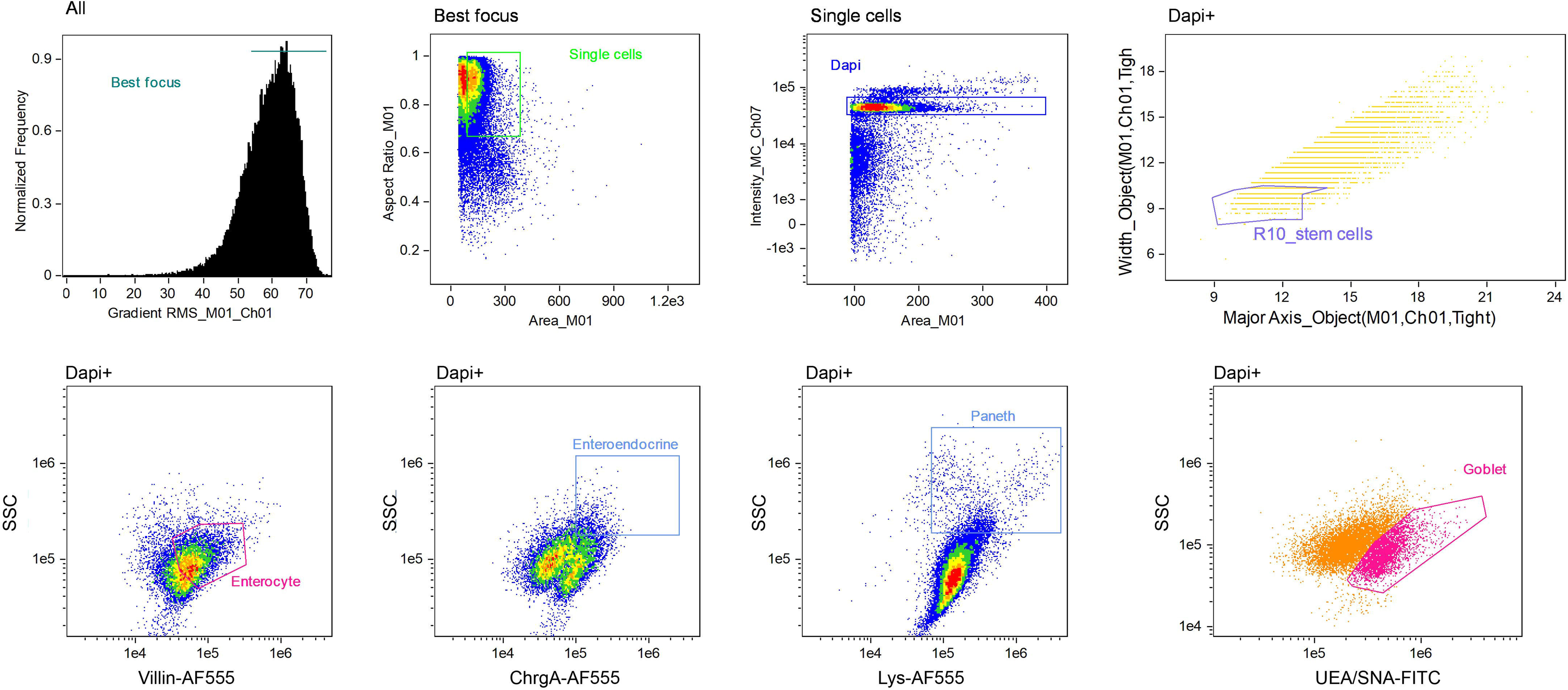
Gating strategy ImageStream. Gates were set up to select for single, in-focus cells using the guided analysis tools for focused and single cell analysis from IDEAS. Specifically, normalised frequency versus Gradient RMS on BF for all events was used to reject out-of-focus events and draw the Best focus gate. On the Best focus population, Aspect ratio versus Area for BF was used to exclude beads and select for single cell events (Single cells gate). “Single cells” were further gated based on nuclear stain to select for optimum nuclear staining by looking at Intensity for emission channel (Ch) 7 versus Area for BF (Dapi gate). All further analysis to quantify epithelial cell populations was performed using “Dapi gate” as initial population. Intensity Ch6 (SSC) versus Ch3 (AF555) or Intensity Ch2 (FITC) was used for determining enterocyte, enteroendocrine, Paneth and goblet cell populations. To measure the stem cell population, the Features Finder tool from IDEAS was used, with Width Object versus Major Axis Object on BF used for gating.

## REFERENCES

Al Alam, D., Sala, F.G., Baptista, S., Galzote, R., Danopoulos, S., Tiozzo, C., Gage, P., Grikscheit, T., Warburton, D., Frey, M.R., Bellusci, S., 2012. FGF9-Pitx2-FGF10 signaling controls cecal formation in mice. Dev Biol 369, 340–348.

Artis, D., Grencis, R.K., 2008. The intestinal epithelium: sensors to effectors in nematode infection. Mucosal immunology 1, 252–264.

Backhed, F., Ley, R.E., Sonnenburg, J.L., Peterson, D.A., Gordon, J.I., 2005. Host-bacterial mutualism in the human intestine. Science 307, 1915–1920.

Bancroft, A.J., Levy, C.W., Jowitt, T.A., Hayes, K.S., Thompson, S., McKenzie, E.A., Ball, M.D., Dubaissi, E., France, A.P., Bellina, B., Sharpe, C., Mironov, A., Brown, S.L., Cook, P.C., A, S.M., Thornton, D.J., Grencis, R.K., 2019. The major secreted protein of the whipworm parasite tethers to matrix and inhibits interleukin-13 function. Nat Commun 10, 2344.

Barker, N., 2014. Adult intestinal stem cells: critical drivers of epithelial homeostasis and regeneration. Nat Rev Mol Cell Biol 15, 19–33.

Barker, N., Huch, M., Kujala, P., van de Wetering, M., Snippert, H.J., van Es, J.H., Sato, T., Stange, D.E., Begthel, H., van den Born, M., Danenberg, E., van den Brink, S., Korving, J., Abo, A., Peters, P.J., Wright, N., Poulsom, R., Clevers, H., 2010. Lgr5(+ve) stem cells drive self-renewal in the stomach and build long-lived gastric units in vitro. Cell Stem Cell 6, 25–36.

Barrila, J., Crabbe, A., Yang, J., Franco, K., Nydam, S.D., Forsyth, R.J., Davis, R.R., Gangaraju, S., Ott, C.M., Coyne, C.B., Bissell, M.J., Nickerson, C.A., 2018. Modeling Host-Pathogen Interactions in the Context of the Microenvironment: Three-Dimensional Cell Culture Comes of Age. Infect Immun 86.

Barthel, M., Hapfelmeier, S., Quintanilla-Martinez, L., Kremer, M., Rohde, M., Hogardt, M., Pfeffer, K., Russmann, H., Hardt, W.D., 2003. Pretreatment of mice with streptomycin provides a Salmonella enterica serovar Typhimurium colitis model that allows analysis of both pathogen and host. Infect Immun 71, 2839–2858.

Bray, N.L., Pimentel, H., Melsted, P., Pachter, L., 2016. Near-optimal probabilistic RNA-seq quantification. Nat Biotechnol 34, 525–527.

Breuer, K., Foroushani, A.K., Laird, M.R., Chen, C., Sribnaia, A., Lo, R., Winsor, G.L., Hancock, R.E., Brinkman, F.S., Lynn, D.J., 2013. InnateDB: systems biology of innate immunity and beyond--recent updates and continuing curation. Nucleic acids research 41, D1228–1233.

Buck, A.H., Coakley, G., Simbari, F., McSorley, H.J., Quintana, J.F., Le Bihan, T., Kumar, S., Abreu-Goodger, C., Lear, M., Harcus, Y., Ceroni, A., Babayan, S.A., Blaxter, M., Ivens, A., Maizels, R.M., 2014. Exosomes secreted by nematode parasites transfer small RNAs to mammalian cells and modulate innate immunity. Nat Commun 5, 5488.

Burns, R.C., Fairbanks, T.J., Sala, F., De Langhe, S., Mailleux, A., Thiery, J.P., Dickson, C., Itoh, N., Warburton, D., Anderson, K.D., Bellusci, S., 2004. Requirement for fibroblast growth factor 10 or fibroblast growth factor receptor 2-IIIb signaling for cecal development in mouse. Dev Biol 265, 61–74.

Chandra, L., Borcherding, D.C., Kingsbury, D., Atherly, T., Ambrosini, Y.M., Bourgois-Mochel, A., Yuan, W., Kimber, M., Qi, Y., Wang, Q., Wannemuehler, M., Ellinwood, N.M., Snella, E., Martin, M., Skala, M., Meyerholz, D., Estes, M., Fernandez-Zapico, M.E., Jergens, A.E., Mochel, J.P., Allenspach, K., 2019. Derivation of adult canine intestinal organoids for translational research in gastroenterology. BMC Biol 17, 33.

Coakley, G., McCaskill, J.L., Borger, J.G., Simbari, F., Robertson, E., Millar, M., Harcus, Y., McSorley, H.J., Maizels, R.M., Buck, A.H., 2017. Extracellular Vesicles from a Helminth Parasite Suppress Macrophage Activation and Constitute an Effective Vaccine for Protective Immunity. Cell Rep 19, 1545–1557.

Collins, J.W., Keeney, K.M., Crepin, V.F., Rathinam, V.A., Fitzgerald, K.A., Finlay, B.B., Frankel, G., 2014. Citrobacter rodentium: infection, inflammation and the microbiota. Nat Rev Microbiol 12, 612–623.

Connor, L.M., Tang, S.C., Cognard, E., Ochiai, S., Hilligan, K.L., Old, S.I., Pellefigues, C., White, R.F., Patel, D., Smith, A.A., Eccles, D.A., Lamiable, O., McConnell, M.J., Ronchese, F., 2017. Th2 responses are primed by skin dendritic cells with distinct transcriptional profiles. J Exp Med 214, 125–142.

Date, S., Sato, T., 2015. Mini-gut organoids: reconstitution of the stem cell niche. Annu Rev Cell Dev Biol 31, 269–289.

Duque-Correa, M.A., Maizels, R.M., Grencis, R.K., Berriman, M., 2019. Organoids - New Models for Host-Helminth Interactions. Trends Parasitol.

Duque-Correa, M.A., Maizels, R.M., Grencis, R.K., Berriman, M., 2020. Organoids - New Models for Host-Helminth Interactions. Trends Parasitol 36, 170–181.

Eckburg, P.B., Bik, E.M., Bernstein, C.N., Purdom, E., Dethlefsen, L., Sargent, M., Gill, S.R., Nelson, K.E., Relman, D.A., 2005. Diversity of the human intestinal microbial flora. Science 308, 1635–1638.

Eichenberger, R.M., Ryan, S., Jones, L., Buitrago, G., Polster, R., Montes de Oca, M., Zuvelek, J., Giacomin, P.R., Dent, L.A., Engwerda, C.R., Field, M.A., Sotillo, J., Loukas, A., 2018a. Hookworm Secreted Extracellular Vesicles Interact With Host Cells and Prevent Inducible Colitis in Mice. Front Immunol 9, 850.

Eichenberger, R.M., Talukder, M.H., Field, M.A., Wangchuk, P., Giacomin, P., Loukas, A., Sotillo, J., 2018b. Characterization of Trichuris muris secreted proteins and extracellular vesicles provides new insights into host-parasite communication. J Extracell Vesicles 7, 1428004.

Fahlgren, A., Avican, K., Westermark, L., Nordfelth, R., Fallman, M., 2014. Colonization of cecum is important for development of persistent infection by Yersinia pseudotuberculosis. Infect Immun 82, 3471–3482.

Fatehullah, A., Tan, S.H., Barker, N., 2016. Organoids as an in vitro model of human development and disease. Nat Cell Biol 18, 246–254.

Grencis, R.K., 2015. Immunity to helminths: resistance, regulation, and susceptibility to gastrointestinal nematodes. Annu Rev Immunol 33, 201–225.

Hansen, E.P., Kringel, H., Williams, A.R., Nejsum, P., 2015. Secretion of RNA-Containing Extracellular Vesicles by the Porcine Whipworm, Trichuris suis. J Parasitol 101, 336–340.

Hiemstra, I.H., Klaver, E.J., Vrijland, K., Kringel, H., Andreasen, A., Bouma, G., Kraal, G., van Die, I., den Haan, J.M., 2014. Excreted/secreted Trichuris suis products reduce barrier function and suppress inflammatory cytokine production of intestinal epithelial cells. Mol Immunol 60, 1–7.

Houpt, E.R., Glembocki, D.J., Obrig, T.G., Moskaluk, C.A., Lockhart, L.A., Wright, R.L., Seaner, R.M., Keepers, T.R., Wilkins, T.D., Petri, W.A., Jr., 2002. The mouse model of amebic colitis reveals mouse strain susceptibility to infection and exacerbation of disease by CD4+ T cells. J Immunol 169, 4496–4503.

James, K.R., Gomes, T., Elmentaite, R., Kumar, N., Gulliver, E.L., King, H.W., Stares, M.D., Bareham, B.R., Ferdinand, J.R., Petrova, V.N., Polanski, K., Forster, S.C., Jarvis, L.B., Suchanek, O., Howlett, S., James, L.K., Jones, J.L., Meyer, K.B., Clatworthy, M.R., Saeb-Parsy, K., Lawley, T.D., Teichmann, S.A., 2020. Distinct microbial and immune niches of the human colon. Nat Immunol 21, 343–353.

Klaver, E.J., Kuijk, L.M., Laan, L.C., Kringel, H., van Vliet, S.J., Bouma, G., Cummings, R.D., Kraal, G., van Die, I., 2013. Trichuris suis-induced modulation of human dendritic cell function is glycan-mediated. Int J Parasitol 43, 191–200.

Klementowicz, J.E., Travis, M.A., Grencis, R.K., 2012. Trichuris muris: a model of gastrointestinal parasite infection. Seminars in immunopathology 34, 815–828.

Kuijk, L.M., Klaver, E.J., Kooij, G., van der Pol, S.M., Heijnen, P., Bruijns, S.C., Kringel, H., Pinelli, E., Kraal, G., de Vries, H.E., Dijkstra, C.D., Bouma, G., van Die, I., 2012. Soluble helminth products suppress clinical signs in murine experimental autoimmune encephalomyelitis and differentially modulate human dendritic cell activation. Mol Immunol 51, 210–218.

Kuipers, M.E., Hokke, C.H., Smits, H.H., Nolte-’t Hoen, E.N.M., 2018. Pathogen-Derived Extracellular Vesicle-Associated Molecules That Affect the Host Immune System: An Overview. Front Microbiol 9, 2182.

Laan, L.C., Williams, A.R., Stavenhagen, K., Giera, M., Kooij, G., Vlasakov, I., Kalay, H., Kringel, H., Nejsum, P., Thamsborg, S.M., Wuhrer, M., Dijkstra, C.D., Cummings, R.D., van Die, I., 2017. The whipworm (Trichuris suis) secretes prostaglandin E2 to suppress proinflammatory properties in human dendritic cells. FASEB J 31, 719–731.

Lee, A., O’Rourke, J.L., Barrington, P.J., Trust, T.J., 1986. Mucus colonization as a determinant of pathogenicity in intestinal infection by Campylobacter jejuni: a mouse cecal model. Infect Immun 51, 536–546.

Leroux, L.P., Nasr, M., Valanparambil, R., Tam, M., Rosa, B.A., Siciliani, E., Hill, D.E., Zarlenga, D.S., Jaramillo, M., Weinstock, J.V., Geary, T.G., Stevenson, M.M., Urban, J.F., Jr., Mitreva, M., Jardim, A., 2018. Analysis of the Trichuris suis excretory/secretory proteins as a function of life cycle stage and their immunomodulatory properties. Sci Rep 8, 15921.

Li, M., Izpisua Belmonte, J.C., 2019. Organoids - Preclinical Models of Human Disease. N Engl J Med 380, 569–579.

Liu, Y., Olagnier, D., Lin, R., 2016. Host and Viral Modulation of RIG-I-Mediated Antiviral Immunity. Front Immunol 7, 662.

Love, M.I., Huber, W., Anders, S., 2014. Moderated estimation of fold change and dispersion for RNA-seq data with DESeq2. Genome Biol 15, 550.

Luo, X.C., Chen, Z.H., Xue, J.B., Zhao, D.X., Lu, C., Li, Y.H., Li, S.M., Du, Y.W., Liu, Q., Wang, P., Liu, M., Huang, L., 2019. Infection by the parasitic helminth Trichinella spiralis activates a Tas2r-mediated signaling pathway in intestinal tuft cells. Proc Natl Acad Sci U S A 116, 5564–5569.

McFarlane, A.J., McSorley, H.J., Davidson, D.J., Fitch, P.M., Errington, C., Mackenzie, K.J., Gollwitzer, E.S., Johnston, C.J.C., MacDonald, A.S., Edwards, M.R., Harris, N.L., Marsland, B.J., Maizels, R.M., Schwarze, J., 2017. Enteric helminth-induced type I interferon signaling protects against pulmonary virus infection through interaction with the microbiota. J Allergy Clin Immunol 140, 1068–1078 e1066.

McGuckin, M.A., Linden, S.K., Sutton, P., Florin, T.H., 2011. Mucin dynamics and enteric pathogens. Nat Rev Microbiol 9, 265–278.

Miyoshi, H., Stappenbeck, T.S., 2013. In vitro expansion and genetic modification of gastrointestinal stem cells in spheroid culture. Nat Protoc 8, 2471–2482.

Mowat, A.M., Agace, W.W., 2014. Regional specialization within the intestinal immune system. Nature reviews. Immunology 14, 667–685.

Nguyen, T.L., Vieira-Silva, S., Liston, A., Raes, J., 2015. How informative is the mouse for human gut microbiota research? Dis Model Mech 8, 1–16.

Perng, Y.C., Lenschow, D.J., 2018. ISG15 in antiviral immunity and beyond. Nat Rev Microbiol 16, 423–439.

Pongpech, P., Hentges, D.J., Marsh, W.W., Eberle, M.E., 1989. Effect of streptomycin administration on association of enteric pathogens with cecal tissue of mice. Infect Immun 57, 2092–2097.

Reynolds, L.A., Harcus, Y., Smith, K.A., Webb, L.M., Hewitson, J.P., Ross, E.A., Brown, S., Uematsu, S., Akira, S., Gray, D., Gray, M., MacDonald, A.S., Cunningham, A.F., Maizels, R.M., 2014. MyD88 signaling inhibits protective immunity to the gastrointestinal helminth parasite Heligmosomoides polygyrus. J Immunol 193, 2984–2993.

Sato, T., Clevers, H., 2013. Growing self-organizing mini-guts from a single intestinal stem cell: mechanism and applications. Science 340, 1190–1194.

Sato, T., Stange, D.E., Ferrante, M., Vries, R.G., Van Es, J.H., Van den Brink, S., Van Houdt, W.J., Pronk, A., Van Gorp, J., Siersema, P.D., Clevers, H., 2011. Long-term expansion of epithelial organoids from human colon, adenoma, adenocarcinoma, and Barrett’s epithelium. Gastroenterology 141, 1762–1772.

Sato, T., Vries, R.G., Snippert, H.J., van de Wetering, M., Barker, N., Stange, D.E., van Es, J.H., Abo, A., Kujala, P., Peters, P.J., Clevers, H., 2009. Single Lgr5 stem cells build crypt-villus structures in vitro without a mesenchymal niche. Nature 459, 262–265.

Shears, R.K., Bancroft, A.J., Hughes, G.W., Grencis, R.K., Thornton, D.J., 2018a. Extracellular vesicles induce protective immunity against Trichuris muris. Parasite Immunol 40, e12536.

Shears, R.K., Bancroft, A.J., Sharpe, C., Grencis, R.K., Thornton, D.J., 2018b. Vaccination Against Whipworm: Identification of Potential Immunogenic Proteins in Trichuris muris Excretory/Secretory Material. Sci Rep 8, 4508.

Smallwood, T.B., Giacomin, P.R., Loukas, A., Mulvenna, J.P., Clark, R.J., Miles, J.J., 2017. Helminth Immunomodulation in Autoimmune Disease. Front Immunol 8, 453.

Takebe, T., Wells, J.M., 2019. Organoids by design. Science 364, 956–959.

Tilney, L.G., Connelly, P.S., Guild, G.M., Vranich, K.A., Artis, D., 2005. Adaptation of a nematode parasite to living within the mammalian epithelium. J Exp Zool A Comp Exp Biol 303, 927–945.

Tritten, L., Tam, M., Vargas, M., Jardim, A., Stevenson, M.M., Keiser, J., Geary, T.G., 2017. Excretory/secretory products from the gastrointestinal nematode Trichuris muris. Exp Parasitol 178, 30–36.

Varyani, F., Fleming, J.O., Maizels, R.M., 2017. Helminths in the gastrointestinal tract as modulators of immunity and pathology. Am J Physiol Gastrointest Liver Physiol 312, G537–G549.

Webb, L.M., Lundie, R.J., Borger, J.G., Brown, S.L., Connor, L.M., Cartwright, A.N., Dougall, A.M., Wilbers, R.H., Cook, P.C., Jackson-Jones, L.H., Phythian-Adams, A.T., Johansson, C., Davis, D.M., Dewals, B.G., Ronchese, F., MacDonald, A.S., 2017. Type I interferon is required for T helper (Th) 2 induction by dendritic cells. EMBO J 36, 2404–2418.

White, R., Kumar, S., Chow, F.W.N., Robertson, E., Hayes, K.S., Grencis, R.K., Duque-Correa, M.A., Buck, A., 2020. Extracellular vesicles from Heligmosomoides bakeri and Trichuris muris contain distinct small RNAs that could enable niche specificity in the host. International Journal of Parasitology.

Yates, A.D., Achuthan, P., Akanni, W., Allen, J., Allen, J., Alvarez-Jarreta, J., Amode, M.R., Armean, I.M., Azov, A.G., Bennett, R., Bhai, J., Billis, K., Boddu, S., Marugan, J.C., Cummins, C., Davidson, C., Dodiya, K., Fatima, R., Gall, A., Giron, C.G., Gil, L., Grego, T., Haggerty, L., Haskell, E., Hourlier, T., Izuogu, O.G., Janacek, S.H., Juettemann, T., Kay, M., Lavidas, I., Le, T., Lemos, D., Martinez, J.G., Maurel, T., McDowall, M., McMahon, A., Mohanan, S., Moore, B., Nuhn, M., Oheh, D.N., Parker, A., Parton, A., Patricio, M., Sakthivel, M.P., Abdul Salam, A.I., Schmitt, B.M., Schuilenburg, H., Sheppard, D., Sycheva, M., Szuba, M., Taylor, K., Thormann, A., Threadgold, G., Vullo, A., Walts, B., Winterbottom, A., Zadissa, A., Chakiachvili, M., Flint, B., Frankish, A., Hunt, S.E., G, I.I., Kostadima, M., Langridge, N., Loveland, J.E., Martin, F.J., Morales, J., Mudge, J.M., Muffato, M., Perry, E., Ruffier, M., Trevanion, S.J., Cunningham, F., Howe, K.L., Zerbino, D.R., Flicek, P., 2020. Ensembl 2020. Nucleic acids research 48, D682–D688.

Zaborin, A., Krezalek, M., Hyoju, S., Defazio, J.R., Setia, N., Belogortseva, N., Bindokas, V.P., Guo, Q., Zaborina, O., Alverdy, J.C., 2017. Critical role of microbiota within cecal crypts on the regenerative capacity of the intestinal epithelium following surgical stress. Am J Physiol Gastrointest Liver Physiol 312, G112–G122.

Zhang, X., Stappenbeck, T.S., White, A.C., Lavine, K.J., Gordon, J.I., Ornitz, D.M., 2006. Reciprocal epithelial-mesenchymal FGF signaling is required for cecal development. Development 133, 173–180.

